# Anticipated Loss of Action Consequences Disrupts Motor Execution in Skilled Basketball Shooting

**DOI:** 10.64898/2026.05.13.722224

**Authors:** Aya Nakao, Norimasa Yamada, Tsubasa Wakatsuki

## Abstract

Internal forward models predict the sensory consequences of motor commands; however, whether the anticipated availability of post-action feedback contributes to the precision of the action itself remains unknown. We manipulated the predictability of post-release visual occlusion in skilled basketball players. Participants performed three-point shots while wearing liquid-crystal shutter goggles. The study tested three conditions: a no-occlusion baseline, certain-occlusion condition in which players knew that their vision would be occluded at ball release in every trial, and random-occlusion condition in which they could not predict whether an occlusion would occur. Shooting accuracy declined in the certain-occlusion condition relative to the no-occlusion condition (49.2% vs 41.7%). The random-occlusion condition did not differ from the baseline (46.1%). Within the random condition, the accuracy in occluded trials were virtually identical to that in non-occluded trials (46.6% vs 46.2%), even though the immediate visual occlusion was the same as in the certain-occlusion condition. These results demonstrate that it is not the absence of post-action information per se that disrupts motor execution, but the prior certainty that action consequences will be unavailable. We interpret this finding as a prospective influence of anticipated consequence loss, whereby motor execution depends on whether the prediction–outcome loop remains closable.

## Introduction

Skilled motor performance relies on the ability of the brain to predict the sensory consequences of its own motor commands. Internal forward models generate such predictions before and during movement, enabling the motor system to compare expected and actual feedback and correct ongoing actions in real time [1,2]. This predictive capacity has traditionally been considered the central mechanism by which the motor system maintains accuracy. A well-tuned forward model allows errors to be detected and corrected before task performance is compromised.

A complementary perspective stems from the ideomotor theory, which holds that actions are selected and controlled by anticipating their sensory consequences [3,4; for a review, see 5]. Within this framework, anticipated effects are not mere by-products of action planning, but are constitutive of it. Kunde [6] demonstrated that simple key-press responses were facilitated when the expected sensory effect was compatible with the response, suggesting that effect anticipation was woven into the motor command itself. More recently, Karsh et al. [7] showed that the mere presence of a temporally contiguous sensory effect following a response enhances the speed and precision of subsequent motor actions, independent of the effect’s informational content.

Together, these findings suggest that predicted sensory consequences are not merely used for correcting errors during movement but may also contribute to the quality of the motor command itself. However, whether this extends to the precision of skilled ballistic movements has not been tested.

Many skilled sports actions are ballistic. Once a movement is launched, the actor cannot exert further physical control over its outcome. A basketball shot, golf swing, and dart throw all fall into this category. Since online correction after launch is physically impossible, post-release sensory information has traditionally been regarded as irrelevant for the control of the action itself—useful for learning across subsequent trials, but unable to influence the current one. However, Quirmbach and Limanowski [8] showed that the cerebellum and posterior parietal cortex compute the expected sensory outcomes before movement onset. Thus, sensory prediction is engaged not only during movement execution but also during action planning. Although their delayed hand movement task involved movements that could be corrected online, the finding that prediction begins during planning before any feedback is available suggests that this process may also operate in ballistic movements where post-release correction is impossible. If this is true, what functions do these predictions serve when corrections are impossible? One possibility is that they serve only the next trial as feedback for learning. Another possibility is that the anticipation of post-action feedback plays a role in stabilising the execution of the current action itself, a role that, to the best of our knowledge, has not been directly tested.

Research on visual control in basketball shooting has provided detailed insights into how visual information is used before and during the shooting motion. Studies using temporal occlusion have demonstrated that vision of the target during the final phase of the shooting motion is critical for accuracy, whereas early visual information contributes less [9,10]. Similarly, quiet eye literature has established that prolonged final fixation on the target prior to ball release is a robust predictor of shooting success [11; for recent reviews, see 12-14]. A common thread across this literature is its focus on visual information that can plausibly guide the shooting motion. Whether the anticipated availability of visual information after release plays a role in execution accuracy has not been examined, which is consistent with the traditional assumption that post-release information does not affect the current shot. In a parallel but separate line of work, Knowledge of Performance (KP) and Knowledge of Results (KR) have been extensively studied as augmented feedback that shapes motor learning across trials [15,16]. Whether the anticipated availability of KP and KR within a single trial contributes to trial precision remains an open question.

This study addressed this question by manipulating the predictability of post-release visual occlusion in skilled basketball players performing three-point shots. Liquid-crystal shutter goggles, triggered by photoelectric sensors at the moment of ball release, were used to create three conditions: (a) a no-occlusion condition, in which players had full visual access after release; (b) a certain-occlusion condition, in which players were informed in advance that their vision would be occluded immediately upon release in every trial, eliminating both ball trajectory and shot outcome information; and (c) a random-occlusion condition, in which players were told that occlusion might occur in each trial without knowing which. This design allowed us to contrast the two competing hypotheses.

In the traditional view, the role of sensory prediction is restricted to online error correction and cross-trial learning. In a ballistic movement, post-release feedback that cannot be acted upon should be irrelevant to the current execution (Hypothesis A). In this view, neither the actual occurrence of occlusion nor its prior anticipation should affect shooting accuracy. Thus, all three conditions should yield comparable performances. An alternative view motivated by ideomotor theory and recent evidence that anticipated action effects contribute directly to motor performance predicts that the anticipated availability of post-action feedback serves as a stabilising condition for the feedforward command itself (Hypothesis B). In this view, accuracy should decline specifically when players are certain that no post-release information will be available, but not when the availability of feedback remains uncertain. A particularly decisive test is provided by the comparison within the random-occlusion condition between trials in which occlusion actually occurred and trials in which it did not. These trials share the same immediate visual occlusion as certain-occlusion trials, yet differ in the prior anticipation of that deprivation. If the physical absence of feedback drives the effect, both should show reduced accuracy. If prior anticipation drives the effect, neither should show reduced accuracy.

## Method

### Participants

A total of 13 female basketball players (mean age = 19.4 ± 1.2 years; mean height = 161.5 ± 4.4 cm) with a minimum of 8 years of competitive experience participated in the study. All participants were members of university women’s basketball teams. Written informed consent was obtained from all the participants prior to the experiment. This study was approved by the Ethics Review Committee for Research Involving Human Subjects of Nihon Fukushi University (Approval No. 18-37)..

### Apparatus

Shots were taken from the regulation three-point line (6.75 m) directly in front of the basket using a standard basketball and regulation-height hoop (3.05 m) in an indoor gymnasium. Visual occlusion was achieved using liquid-crystal shutter goggles (ToTaL Control System; Translucent Technologies, Toronto, Canada). According to the manufacturer’s specifications, the goggles transition from a transparent to fully occluded state in approximately 7 milliseconds (ms). To time-lock the occlusion to the moment of ball release, photoelectric sensors (E3Z-61; OMRON, Kyoto, Japan) were mounted on two vertical bars positioned on either side of the shooting trajectory, 0.725 m from the release point.

Prior to the main experiment, each participant performed several practice shots so that the height of the sensors could be adjusted to match the trajectory of their release. When the ball passed through the sensor beam shortly after its release, the sensor signal triggered the goggles to switch to an opaque state. This system ensured that the onset of occlusion was determined by the ball release without the experimenter’s intervention. Based on previously reported release velocities for three-point shots in female basketball players (7.82 m/s) [17], the ball typically crossed the sensor approximately 93 ms after release, yielding an estimated latency from ball release to full visual occlusion of approximately 100 ms. However, the precise temporal characteristics of goggle activation (approximately 100 ms post-release) were not central to the experimental logic. What mattered was the structural distinction between the conditions. Therefore, the conclusions did not depend on millisecond-level timing precision. Figure 1A shows the experimental setup and Figure 1B provides a schematic illustration of the goggle occlusion state timeline and temporal delay.

**Figure 1.**
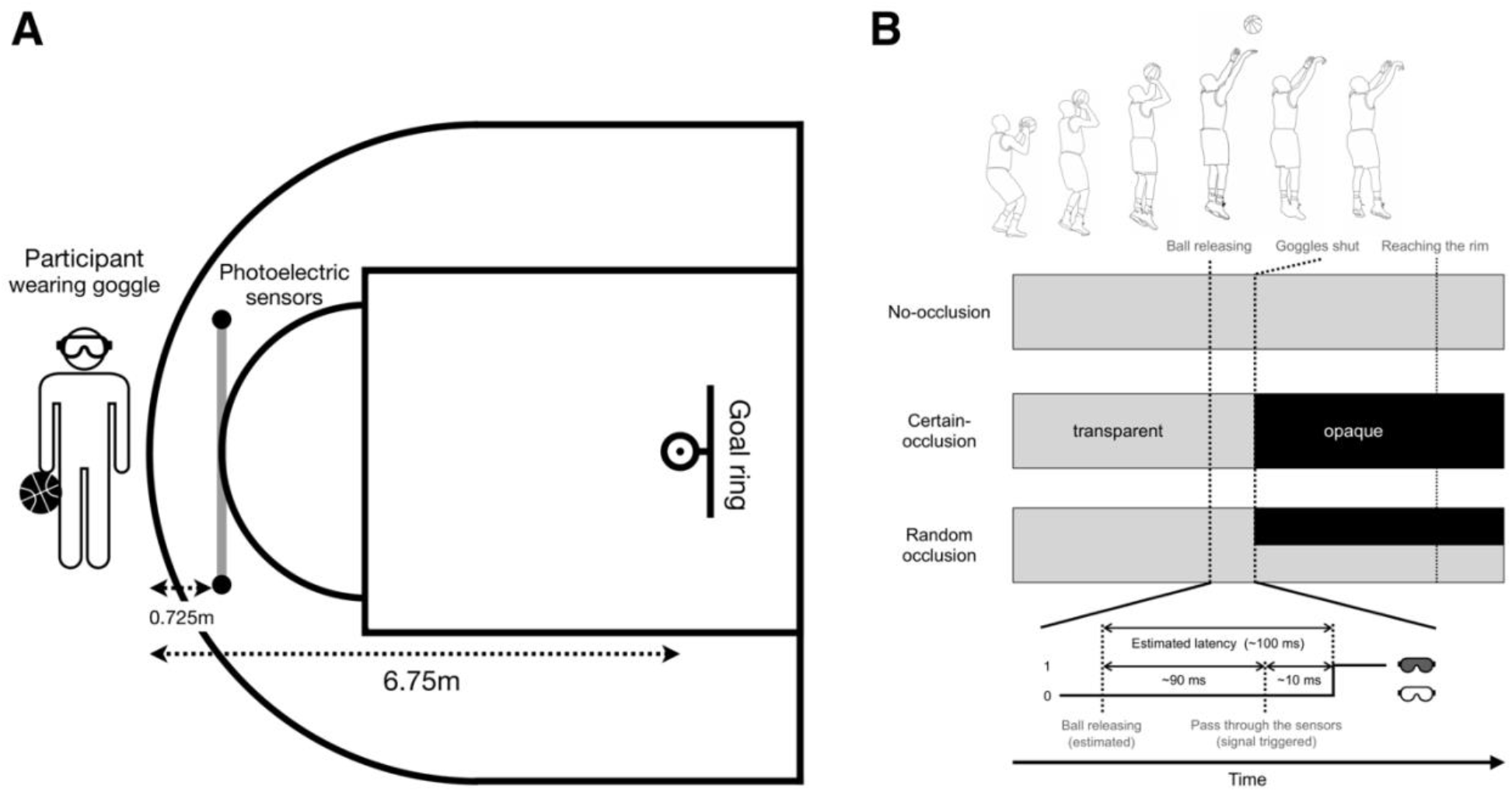
Experimental setup. (A) Top-down view of the shooting arrangement. A participant wearing liquid-crystal shutter goggles performed shots from the three-point line (6.75 m from the goal ring). Two photoelectric sensors were positioned 0.725 m in front of the participant to detect the moment of ball release; when the ball passed through the sensor beam, the signal triggered the goggles to switch from transparent to opaque. (B) Timeline of goggle state across the three conditions. Gray bars indicate the transparent state and black bars indicate the opaque state. In the random-occlusion condition, occluded and non-occluded trials were randomly selected with 50% probability each. Goggle occlusion did not occur precisely at ball release but approximately 100 ms thereafter, reflecting the time it takes for the ball to reach the photoelectric sensor and the technical response limit of the goggles. The manipulation targets the availability of visual consequences of the self-generated ball immediately after release and the certainty of this loss rather than precise millisecond timing.

### Procedure

Before the main experiment, the participants performed unrestricted warm-up shooting until they felt fully prepared. They then put on the shutter goggles and performed additional familiarisation shots until they felt comfortable shooting while wearing the goggles. The participants wore the shutter goggles in all three conditions to ensure that any discomfort or visual restriction caused by the goggles was constant across the conditions.

The participants performed three-point shots under three conditions: no-occlusion, certain-occlusion, and random-occlusion. (1) No-occlusion condition: the goggles remained transparent throughout the trial. The participants had full visual access before, during, and after the shot. (2) Certain-occlusion condition: the participants were informed before the block that their vision would be occluded immediately upon the ball release in every trial. (3) Random-occlusion condition: the participants were told that occlusion might occur in any given trial. Thus, they could not predict in which trials occlusion would occur. Within each 20-trial set of the random-occlusion condition, occluded and non-occluded trials were randomly selected with 50% probability each. Thus, across the 80 trials of the random-occlusion condition, each participant experienced approximately 40 occluded and 40 non-occluded trials. Under both occlusion conditions, the goggles became opaque at the moment of release, eliminating visual access to the flight trajectory of the ball and shot outcomes. Figure 1B shows a conceptual diagram of the time series of the three conditions.

Each condition comprised 80 trials divided into four sets of 20 trials each. The three conditions were always presented in 20-trial blocks and never intermixed within a block so that participants always knew which condition they were performing throughout each block. The order of the three conditions was counterbalanced across participants and sets within participants. Each participant experienced a different sequence of the 12 blocks (3 conditions × 4 sets), and the order of the three conditions was rotated across sets within each participant. Rest periods of sufficient duration were provided between sets to prevent fatigue. A total of 240 shots were performed per participant, yielding 3,120 trials across the sample.

### Data Analysis

The shot outcome (successful or unsuccessful) was recorded for each trial by an experimenter. Shooting accuracy was calculated as the percentage of successful shots per condition for each participant. Frequentist analyses were performed using Mathematica (version 12.2; Wolfram Research, Champaign, IL, USA).

A one-way repeated-measures analysis of variance (ANOVA) was conducted on shooting accuracy across the three conditions, followed by pairwise comparisons with Holm correction [18]. Mauchly’s test was used to assess the sphericity. Partial eta-squared (η^2^p) was reported as a measure of effect size for the omnibus test, and Cohen’s d was reported for all pairwise comparisons. Throughout the Results, Mdiff denotes the mean difference between the two conditions and p adjusted denotes the Holm-corrected p-value.

To test whether performance within the random condition depended on whether occlusion occurred in a given trial, shooting accuracy in occluded and non-occluded trials was compared using a paired-samples t-test. Since this analysis was central to our theoretical interpretation and relied on the absence of an effect, a Bayes factor was additionally computed using the default Jeffreys-Zellner-Siow prior (Cauchy scale r = 0.707) in JASP (version 0.18.3) [19] to quantify the strength of the evidence for the null hypothesis. The Bayes factor was computed only for this critical comparison because it provided the most direct test of whether the physical occurrence of occlusion (rather than its anticipation) influences motor execution.

As an exploratory analysis, we also examined whether accuracy in the random condition was affected by the occlusion status of the preceding trial. Accuracy was calculated separately for trials preceded by an occluded trial versus trials preceded by a non-occluded trial. The first trial of each 20-trial set was excluded from this analysis because it was not preceded by a trial within the same set. The values were compared using paired-sample t-tests. All analyses used an alpha level of .05.

To confirm that the condition effects were not artefacts of aggregating trial-level data to participant-level accuracy scores and to provide direct contrasts between conditions at the trial level, we conducted a supplementary Bayesian mixed-effects logistic regression on the trial-level data (3,120 observations). The model predicted the trial-level shot outcome (success or failure) from the condition as a fixed effect with a by-participant random intercept to account for individual differences in overall shooting accuracy. We used a Bernoulli distribution with a logit link function. The model was fitted using JASP (version 0.18.3; JASP Team, 2024) with default priors. Three Markov Chain Monte Carlo (MCMC) chains were run with 2,000 warm-up iterations and 4,000 post-warm-up samples each. Furthermore, the adapt delta was set to 0.95 to minimize divergent transitions. Planned contrasts on the response scale were computed to directly compare the three conditions: no-occlusion versus certain-occlusion, no-occlusion versus random-occlusion, and certain-occlusion versus random-occlusion. We reported the posterior medians, 95% highest posterior density intervals (HPD), and estimated the marginal means for each condition.

## Results

The descriptive statistics for shooting accuracy across the three conditions are presented in Figure 2A. The mean shooting accuracies were 49.2% (SD = 7.2), 41.7% (SD = 9.9), and 46.1% (SD = 7.7) under the no-occlusion, certain-occlusion, and random-occlusion conditions, respectively. A one-way repeated-measures ANOVA revealed a significant main effect of condition, F (2, 24) = 7.73, p = .003, η^2^p = .392. Mauchly’s test indicated that the assumption of sphericity had not been violated, χ^2^ (2) = 1.59, p = .451 (Greenhouse-Geisser ε = .88). Pairwise comparisons with Holm correction showed that accuracy was significantly lower in the certain-occlusion condition than in the no-occlusion condition (Mdiff = 7.51%, t (12) = 4.22, p adjusted = .004, d = 1.17). The difference between the no-occlusion and random-occlusion conditions did not reach significance (Mdiff = 3.17%, t (12) = 1.88, p adjusted = .084, d = 0.52), nor did the difference between the certain-occlusion and random-occlusion conditions (Mdiff = 4.33%, t (12) = 1.93, p adjusted = .154, d = 0.54).

**Figure 2.**
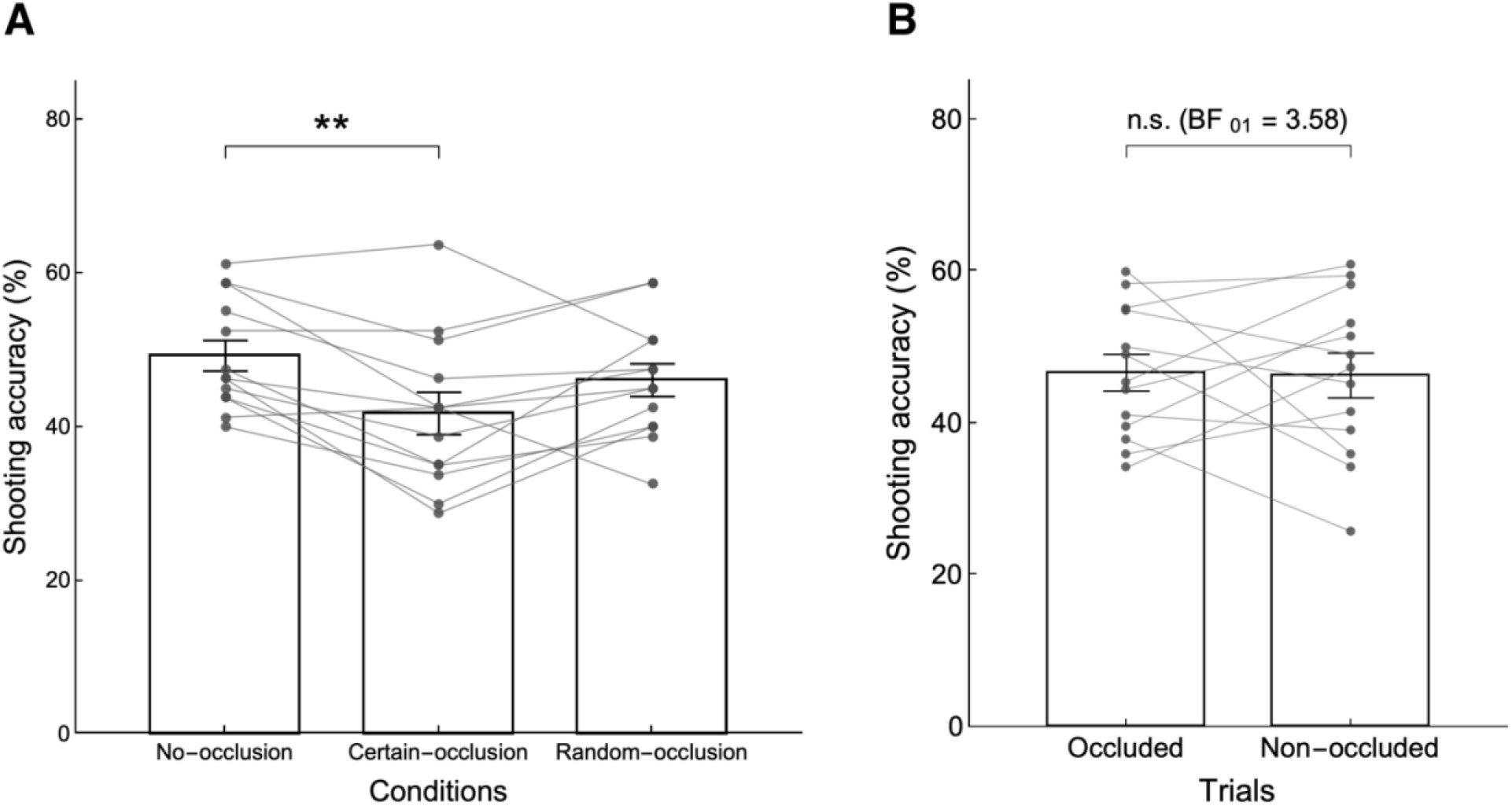
Shooting accuracy. (A) Mean shooting accuracy (%) across the three conditions. Bars represent group means with standard error bars; grey dots show individual participants connected by lines across conditions. **p < .01. (B) Mean shooting accuracy within the random-occlusion condition for occluded and non-occluded trials, with the same conventions as in panel A. ‘n.s.’ denotes a non-significant difference (p = .916); BF_01_ = 3.58 indicates moderate Bayesian evidence in favour of the null hypothesis.

Within the random-occlusion condition, accuracy in trials in which occlusion actually occurred (46.6%, SD = 8.7) was virtually identical to accuracy in trials in which it did not (46.2%, SD = 10.6), Mdiff = 0.35%, t (12) = 0.11, p = .916, d = 0.03, BF_01_ = 3.58 (Figure 2B). The Bayes factor indicated moderate evidence in favour of the null hypothesis [20]. This suggests that the actual occurrence of occlusion within the random condition had no measurable effect on accuracy. An exploratory carry-over analysis examined whether the accuracy in the random condition was affected by the occlusion status of the preceding trial. The accuracy following an occluded trial (46.1%, SD = 8.2) did not differ significantly from that following a non-occluded trial (46.1%, SD = 9.4), Mdiff = 0.03%, t (12) = 0.01, p = .989, d = 0.004.

To address concerns that aggregating across trials could obscure condition effects and provide direct trial-level comparisons between conditions, we conducted a supplementary Bayesian mixed-effects logistic regression on the trial-level data. The estimated marginal means on the response scale were 0.416 (95% HPD [0.367, 0.464]) for certain-occlusion, 0.492 (95% HPD [0.440, 0.541]) for no-occlusion, and 0.460 (95% HPD [0.409, 0.508]) for random-occlusion, closely matching the participant-level descriptives. Planned contrasts confirmed the pattern observed in the participant-level analyses. The accuracy was higher in the no-occlusion condition than in the certain-occlusion condition (difference = 0.076, 95% HPD [0.030, 0.122]), with the interval excluding zero. In contrast, accuracy in the no-occlusion condition did not differ reliably from the random-occlusion condition (difference = 0.032, 95% HPD [−0.017, 0.080]), with the interval spanning zero. The contrast between the certain-occlusion and random-occlusion conditions was in the expected direction but with the interval marginally including zero (difference = −0.044, 95% HPD [−0.091, 0.003]). All R-hat values were ≤ 1.002, indicating adequate MCMC convergence. These trial-level results directly confirmed the pattern observed in the participant-level analyses. Accuracy declined selectively in the certain-occlusion condition, whereas the random-occlusion condition was not distinguishable from the no-occlusion baseline.

## Discussion

Through an experimental paradigm that manipulated the predictability of post-release visual occlusion in skilled basketball players, the present findings demonstrate that the certainty of post-release consequence loss, rather than consequence loss itself, disrupts skilled motor execution. Shooting accuracy declined significantly only when the players knew in advance that every shot would be followed by complete visual occlusion. When occlusion was unpredictable, the accuracy was not statistically distinguishable from baseline, even in trials in which occlusion actually occurred. Immediate visual occlusion was the same as in the certain-occlusion condition. The experimental manipulation did not hinge on the millisecond-level precision of occlusion timing; rather, it controlled whether visual information about the self-generated ball becomes unavailable immediately after release and, critically, whether this loss of post-release visibility was certain or uncertain across conditions.

This pattern cannot be explained by the absence of post-release information. If possible, the occluded trials within the random condition should have shown the same decrement as in the certain-occlusion condition. Instead, accuracy on occluded and non-occluded trials within the random condition was virtually identical, with Bayesian analysis providing moderate evidence for the null hypothesis (BF_01_ = 3.58). What distinguished the certain-occlusion condition from occluded trials in the random condition was not what happened after release but what players anticipated before it. We propose that this reflects the prospective influence of anticipated consequence loss, in the sense that motor execution depends on whether the prediction–outcome loop remains structurally closable. Table 1 summarises how each condition and trial type maps onto the key theoretical predictors of accuracy. Of these, only reachability, the structural possibility of accessing outcome information such that the prediction–outcome loop remains, in principle, closable, shows a systematic correspondence with the observed pattern.

**Table 1.** Theoretical predictors of shooting accuracy across conditions and trial types.

| Condition/Trial type | Entropy (bits) | Reachability | Anticipation of effect | Actual feedback | Shooting accuracy (%) |
| --- | --- | --- | --- | --- | --- |
| No-occlusion | 0 | Open | Confirmed | Available | 49.2 |
| Certain-occlusion | 0 | Foreclosed | Confirmed (adverse) | Unavailable | 41.7 |
| Random-occlusion | 1 | Open |  |  | 46.1 |
| *on non-occluded* |  |  | Confirmed | Available | 46.2 |
| *on occluded* |  |  | Disconfirmed | Unavailable | 46.6 |

*Note.* Reachability indicates whether the structural possibility of accessing the post-release outcome information is preserved (open) or precluded in advance (foreclosed). Entropy represents trial-level information-theoretic uncertainty regarding the occurrence of visual occlusion. The anticipation of the effect refers to whether the actor's prior expectation of post-release visual information was confirmed (matched the actual outcome) or unconfirmed (did not match). Confirmed (adverse) in the certain-occlusion condition indicated that the prior expectation was confirmed, but the expectation itself was an unfavourable outcome (loss of visual feedback). Actual feedback refers to whether the post-release visual information was physically available during a given trial. The bold row shows the overall values for the random-occlusion condition, and the indented italic rows show the values for each trial type within that condition. Shooting accuracy systematically corresponds to reachability but not to entropy, anticipation, or the presence of actual feedback.

### Action-effect Anticipation and Motor Execution

This finding resonates with the central tenet of ideomotor theory, namely, that actions are selected and controlled through the anticipation of their sensory consequences [3,4; for a review, see 5]. In the ideomotor framework, anticipated effects are not merely byproducts of action planning; they are constitutive of it. Kunde [6] demonstrated that even simple key-press responses were facilitated when the expected sensory effect was compatible with the response, suggesting that effect anticipation was woven into the motor command itself. The present results extend this principle from action selection to action precision: when the anticipated effects of a skilled action are pre-emptively nullified, that is, when the actor knows that no consequence will be observable, the temporal and spatial stability of the motor output deteriorates.

Recent studies have provided further evidence that action effects directly contribute to motor performance. Karsh et al. [7] showed that the mere presence of a temporally contiguous sensory effect following a response enhances the speed and precision of subsequent motor actions, independent of its informational content. Their interpretation, grounded in the Control-Based Response Selection framework [21], holds that the confirmed sensorimotor predictions carry intrinsic value for the motor system. Our findings complement this perspective by demonstrating the converse: when the confirmation of sensorimotor predictions is guaranteed to be impossible, motor precision suffers.

### Forward Models and the Structural Availability of Outcome Information

These results are consistent with the framework of internal forward models that predict the sensory consequences of motor commands [1,8]. In the standard account, these predictions serve as online corrections during movement and are updated through comparisons with actual sensory feedback [2]. For ballistic movements such as basketball shots, online correction after release is impossible, yet the forward model presumably still generates a prediction of the post-release trajectory. More broadly, this perspective suggests that from the standpoint of the motor system, action and its anticipated consequences may not be separable temporal events but rather form a single integrated structure whose stability depends on the potential closure of the prediction– outcome loop. When the system is informed in advance that the consequence portion of this unit will be unavailable, the integrity of the entire representation, including the motor commands preceding the consequence, may be compromised.

The present findings indicate that it is not the physical absence of post-release visual information per se that degrades performance but the prior certainty that such information will be unavailable. Critically, in the random-occlusion condition, trials with actual occlusions did not differ from non-occluded trials, ruling out the physical absence of feedback as the primary driver. We interpreted this pattern in terms of the structural availability of the outcome information (see Table 1, where reachability is the only predictor that systematically tracks the pattern of accuracy). Rather than being a point event, an action can be described as a process organised with respect to whether its outcome remains reachable within the task structure. In principle, the motor output is stabilised when the prediction–outcome loop remains closable.

When this closure is precluded in advance, the motor system cannot maintain a stable feedforward command, resulting in degraded execution precision. This account is consistent with ideomotor theory, which posits that actions are linked to their anticipated effects. However, the present data further suggest that it is not merely the anticipation of effects but also the structural possibility of accessing those effects that contributes to execution precision. Importantly, this interpretation is intended as a functional description rather than a claim about a specific neural implementation.

The information eliminated by occlusion included both the ball’s flight trajectory (KP) and shot outcome (KR). Under normal conditions, both forms of feedback are conventionally understood as learning signals guiding subsequent performance [15,16]. The present findings raise the possibility that anticipated availability also plays a role in stabilising execution in the current trial, a function that is distinct from and logically prior to their role in learning. This distinction has implications for how augmented feedback is conceptualised in motor learning theory: feedback may serve not only to update internal models retrospectively but also to sustain the precision of the motor output that generates the actions being evaluated.

An important consideration is the nature of the post-release visual feedback in skilled basketball shooting. Consistent with the view that internal models linking actions to their sensory consequences are progressively shaped by practice [1,22], expert players have likely formed strong perception–motor couplings through extensive practice, in which the visual consequences of release are not external information that arrives after the action but constitutive elements of the integrated motor representation itself. From this perspective, what must remain reachable is not merely abstract outcome information but the learned perceptual–motor coupling embedded within the act. The prediction–outcome loop itself is a learned structural feature of skilled action rather than an external feedback channel. The certain-occlusion condition forecloses this internalised structure in advance.

This view may also invite a tentative theoretical reflection. Traditional accounts of motor control distinguish between closed-loop control, in which sensory feedback is used online to correct ongoing movement, and open-loop control, which is characteristic of ballistic actions in which no online correction is possible [23,24]. However, the present pattern suggests a dimension beyond this distinction. As discussed, previously, in skilled ballistic action, post-release feedback is not an external information channel that the system can elect to use but a learned structural feature embedded within the motor representation itself. In this view, what disrupts execution in the certain-occlusion condition is therefore not the absence of online correction (impossible in any condition here) or the absence of feedback per se (matched between certain-occlusion and occluded random-occlusion trials), but the prior foreclosure of this internalised prediction– outcome relation. A state in which this internalised relationship remains structurally available for closure may be tentatively termed a closable-loop. Under this interpretation, motor stability in skilled ballistic action may depend not on the use or non-use of feedback during execution but on whether the embedded prediction–outcome loop remains, in principle, closable. Further research across tasks, modalities, and species is required to assess the generalisability of this proposal.

### The Paradox of Certainty-dependent Disruption

The pattern of results is difficult to reconcile with accounts in which unpredictability imposes costs on motor performance. As shown in Table 1, the random-occlusion condition carried the greatest uncertainty from an information-theoretic perspective (1 bit of entropy regarding the upcoming visual state), whereas both the no-occlusion and certain-occlusion conditions were fully predictable (0 bits). Several classes of accounts would predict that the performance should decline with increasing entropy. Classical studies of temporal preparation have shown that reaction times increase when the timing of an upcoming stimulus becomes uncertain [25]. This principle has been extended to uncertainty regarding whether the stimulus will occur at all [26; for a review, see 27]. Similarly, Bayesian accounts of motor control predict that uncertainty about upcoming events destabilises motor output [28]. Models based on limited attentional resources [29] predict that monitoring an uncertain environment consumes capacity that would otherwise support execution, thereby degrading performance.

Finally, a long tradition in the psychology of performance under pressure holds that unpredictability elicits arousal and anxiety, which can narrow attention and disrupt skilled actions [30,31]. Each framework predicts the greatest decrement under the random-occlusion condition. However, the present data show the opposite asymmetry; performance was preserved under maximum entropy and disrupted under minimum entropy, with an adverse expectation.

This pattern contradicts explanations framed in terms of unpredictability, cognitive load from uncertainty, attentional resource competition, or arousal induced by an uncertain environment. What the certain-occlusion condition uniquely provides is not uncertainty but its opposite: a confident prior expectation that post-release feedback will be unavailable. This reinforces the interpretation that the critical variable is not uncertainty itself but the prior certainty of consequence loss (see Table 1, where entropy is decoupled from accuracy). In the framework, this prior certainty corresponds to the structural foreclosure of outcome reachability. The certain-occlusion condition uniquely removes the structural possibility of accessing post-release feedback, whereas the random-occlusion condition preserves this possibility, regardless of the trial-level outcome.

### Limitations and Future Directions

This study had several limitations. First, auditory information (e.g. the sound of the ball contacting the rim or passing through the net) was not controlled, and participants may have inferred shot outcomes from auditory cues in some trials. However, this auditory information was equally available across all three conditions and therefore cannot account for the observed differences between the conditions. Moreover, the fact that accuracy declined in the certain-occlusion condition despite the availability of auditory KR suggests that the anticipated loss of visual consequence information was the primary driver of this effect.

Second, the sample size (N = 13) was modest. A sensitivity analysis conducted with G*Power (version 3.1) [32] for a within-factors repeated-measures ANOVA (α = .05, power [1 − β] = .80, number of measurements = 3, correlation among repeated measures = .50, nonsphericity correction ε = 1) indicated that this sample size provided adequate power to detect effects of f = 0.374 or larger. The observed effect size was large (η^2^p = .392, corresponding to f = 0.803), substantially exceeding this threshold. Each participant completed 240 trials (3,120 total). The critical within-condition comparison in the random condition yielded p = .916 with moderate Bayesian evidence for the null (BF_01_ = 3.58), suggesting a robust pattern.

Finally, the present study only measured shooting accuracy, which limits our ability to specify which aspects of the shooting action were altered. A particularly informative follow-up would involve electromyographic recording of the major muscles involved in shooting. Under our interpretation, the certain-occlusion condition should be associated with increased trial-to-trial variability in muscle activation, consistent with a destabilised feedforward command [33], and possibly with greater agonist–antagonist co-contraction, reflecting a compensatory stiffening strategy employed when precise motor control is required [34]. Critically, such changes should be observable before ball release, which is consistent with the evidence that movement variability is largely determined during motor preparation [35]. This provides direct evidence that the anticipation of post-release feedback loss affects motor preparation. Under the same interpretation, the within-random null findings should be extended to these muscular measures. While a simple attentional or motivational account cannot easily explain the null difference between occluded and non-occluded trials within the random condition, fully disentangling behavioural and mechanistic interpretations will require future work combining behavioural measures with electromyography.

## Conclusion

The present study provides evidence that skilled motor performance depends not only on the sensory information available during action preparation and execution but also on the anticipated availability of information that will arrive only after the action is complete. Prior certainty that visual action consequences will be unavailable undermines precision, whereas mere uncertainty about their availability does not. This prospective influence, in which the expected status of a future event shapes the quality of a preceding action, opens a new perspective on the relationship between action and consequence in motor control. This finding suggests that the motor system treats execution and its anticipated consequences as an integrated unit whose stability depends on the structural possibility of closing the prediction–outcome loop.

